# A novel approach to partitioning evapotranspiration into evaporation and transpiration in flooded ecosystems

**DOI:** 10.1101/2021.04.06.438244

**Authors:** Elke Eichelmann, Mauricio C. Mantoani, Samuel D. Chamberlain, Kyle S. Hemes, Patricia Y. Oikawa, Daphne Szutu, Alex Valach, Joseph Verfaillie, Dennis D. Baldocchi

## Abstract

Reliable partitioning of micrometeorologically measured evapotranspiration (ET) into evaporation (E) and transpiration (T) would greatly enhance our understanding of the water cycle and its response to climate change. While some methods on ET partitioning have been developed, their underlying assumptions make them difficult to apply more generally, especially in sites with large contributions of E. Here, we report a novel ET partitioning method using Artificial Neural Networks (ANN) in combination with a range of environmental input variables to predict daytime E from nighttime ET measurements. The study uses eddy covariance data from four restored wetlands in the Sacramento-San Joaquin Delta, California, USA, as well as leaf-level T data for validation. The four wetlands vary in structure from some with large areas of open water and little vegetation to very densely vegetated wetlands, representing a range of ET conditions. The ANNs were built with increasing complexity by adding the input variable that resulted in the next highest average value of model testing R^2^ across all sites. The order of variable inclusion (and importance) was: vapor pressure deficit (VPD) > gap-filled sensible heat flux (H_gf) > air temperature (T_air_) > friction velocity (u_∗_) > other variables. Overall, 36 ANNs were analyzed. The model using VPD, H_gf, T_air_, and u_∗_ (F11), showed an average testing R^2^ value across all sites of 0.853. In comparison with the model that included all 10 variables (F36), F11 generally performed better during validation with independent data. In comparison to other methods described in the literature, the ANN method generated more consistent T/ET partitioning results especially for more complex sites with large E contributions. Our method improves the understanding of T/ET partitioning. While it may be particularly suited to flooded ecosystems, it can also improve T/ET partitioning in other systems, increasing our knowledge of the global water cycle.

## 1 Introduction^1^

Evapotranspiration^2^ (ET) is the combined water loss from terrestrial ecosystems via transpiration (T), i.e., water lost by plants during the process of carbon assimilation, and evaporation (E), i.e., water lost via direct evaporation of soil and surface water. Through these processes, ET adds on the order of 65 to 75 thousand km^3^ of water to the atmosphere every year (Oki & Kanae, 2006; Trenberth, Fasullo, & Kiehl, 2009; Jung et al., 2018) and constitutes an important component of the terrestrial water cycle. Despite its importance to the global water cycle, ET is currently poorly constrained in global land surface models (LSM), and it is unclear whether ET will increase or decrease with climate change which creates large uncertainties in climate predictions (Brutsaert & Parlange, 1998; Zeng et al., 2018). This is partly because E and T have different drivers and mechanisms. Thus, improving our understanding of the relative contribution of E and T to ET will improve our ability to predict how the water cycle will evolve with climate change (Stoy et al., 2019).

Assessments of E and T fluxes at an ecosystem scale (i.e., 100 m to km) have been attempted using a variety of methods (Stoy et al., 2019). While some methods attempt to determine E and T components by direct measurements (e.g., measurement of soil evaporation, sap-flux measurements for transpiration), these are often time and labor intensive and present significant challenges upscaling results to ecosystem level (Wilson et al., 2001). Micrometeorological methods, such as eddy covariance (EC), are well-established methods that assess biosphere-atmosphere fluxes of trace gases at the ecosystem scale (Baldocchi et al., 1988). With EC (see Fluxnet.org, 2021) continuous measurements of ecosystem trace gas fluxes such as water vapor can be made on time scales from individual half hours to years (Baldocchi, 2003). However, it can generally only provide direct measurements of the net biosphere-atmosphere flux above the plant canopy. In the case of water vapor fluxes, this includes the net flux of E and T combined. The ability to partition micrometeorologically measured ET fluxes into E and T components would greatly improve our understanding of the pathways by which ecosystems use water, including how E and T components change on different timescales and with changing climatic conditions, as well as the impact of site-specific characteristics like vegetation cover heterogeneity (Eichelmann et al., 2018).

While there are several well tested and established methods to partition net ecosystem CO_2_ fluxes into its components of gross primary production and ecosystem respiration (Baldocchi, 2003; Reichstein et al., 2005; Desai et al., 2008), less work has been done on partitioning ET fluxes (Stoy et al., 2019). Stoy et al. (2019) provide a review of the most common methods for determining E and T fluxes at ecosystem level. Most methods proposed for partitioning micrometeorologically measured ET fluxes use the intrinsic relationship between CO_2_ uptake and transpirational water loss, linked through stomatal exchange at the plant level, to estimate ecosystem T (e.g., Scanlon and Sahu, 2008; Zhou et al., 2016; Scott and Biederman, 2017; Nelson et al., 2018; Li et al., 2019). Scott and Biederman (2017) proposed a method to partition long-term ET measurements into E and T. Their method provides multi-year averages of partitioning on a weekly to yearly timescale. However, it requires datasets of multiple year lengths with high interannual consistency in seasonal ecosystem ET behavior. Furthermore, it is unclear if this method provides reliable results in systems that have a large contribution of E or large interannual variation in ecosystem water exchange behavior.

Similarly, the partitioning method proposed by Scanlon and Sahu (2008), Scanlon and Kustas (2010), and Skaggs et al. (2018), uses the correlation between the high frequency fluctuation of water vapor and CO_2_ concentrations to determine the stomatal and non-stomatal mediated components of the net water and CO_2_ fluxes. However, this method relies on the knowledge of water use efficiency (WUE), which is the ratio of carbon uptake through photosynthesis to water loss through T, at the plant or leaf-level. Since information on WUE is not always readily available at the temporal scale required for this method, and because WUE can change over time with successional age and environmental factors like CO_2_ fertilization, it restricts the wider use of this method. Another method based on the relationship between CO_2_ uptake and T proposed by Zhou et al. (2016) to partition ET data from EC measurements works with the underlying assumption that there will be periods for which E is zero and T/ET approaches one. Similarly, the method proposed by Nelson et al. (2018) assumes that the ecosystem will be dominated by T for some time periods. While such methods are an advancement on T/ET partitioning, there is space for other new approaches particularly if they do not need specialized data or costly equipment to increase the wider use and applicability of such techniques.

Ecosystems with large contributions of E, where total ET is not always dominated by T and which have complex interrelationships between ecosystem productivity, E, and T, might violate some or all of the underlying assumptions necessary for partitioning methods based on the relationship between CO_2_ uptake and water loss to work (Stoy et al., 2019). This is the case for wetlands, where the contribution of E-T is altered significantly by structural factors such as areas of open water, as well as environmental factors, for instance, diurnal fluctuations in air or water temperature and water table (Drexler et al., 2004; Goulden et al., 2007; Eichelmann et al., 2018). In addition, the before-mentioned methods only work when the ecosystem CO_2_ flux is known in conjunction with ET. Although this is often the case for EC measurements, there are other micrometeorological methods that provide measurements of ET without measuring CO_2_ fluxes. Consequently, a partitioning method that does not rely on knowledge of CO_2_ flux and assumptions of carbon-water flux correlations would greatly enhance our ability to partition T/ET in a diversity of settings.

Methods applied to partition CO_2_ fluxes usually use relationships of environmental drivers with the individual flux components determined from time periods where only one flux component is present and extrapolate these to the other periods (Reichstein et al., 2005; Desai et al., 2008). Many methods (e.g., Barr et al., 2004; Reichstein et al., 2005) use relationships between temperature and ecosystem respiration based on nighttime fluxes, when CO_2_ uptake is zero, and extrapolate these to calculate daytime ecosystem respiration. The gross CO_2_ uptake component is then determined as the difference between the net flux and the estimated daytime ecosystem respiration. While this method works well for carbon flux partitioning, where the primary driver of ecosystem respiration is considered to be temperature, it can face limitations in the case of water fluxes where nighttime fluxes are often very small and the drivers of E and T are complex. However, it has been shown that nighttime T from plants is usually very small in many ecosystems (Caird et al., 2006; Dawson et al., 2007). Thus, for non-water limited systems with large contributions of E, such as wetlands, we can approximate nighttime water fluxes as exclusively E.

A newer approach used to partition net ecosystem carbon fluxes into the individual components of gross primary production and ecosystem respiration uses Artificial Neural Networks (ANN) (Papale & Valentini, 2003; Desai et al., 2008; Tramontana et al., 2020). Although the use of ANNs could also be directed at T/ET partitioning, the application of this technique has not been done yet and needs further exploration. Since machine learning methods can resolve complex, nonlinear relationships between environmental drivers and flux variables (Tramontana et al., 2020), ANNs are a promising approach to partition T/ET in ecosystems where existing ET partitioning methods face limitations, such as wetlands and river deltas.

The Sacramento-San Joaquin River Delta (hereafter, the Delta) plays an essential role in the water supply of the state of California, USA. The Delta supplies the majority of freshwater to large metropolises in Southern California and provides water for irrigation of crops in the Central Valley (Deverel & Rojstaczer, 1996). Historically, the Delta’s peat soils were flooded with large areas of freshwater marsh, but the majority of the Delta land area is now actively drained and cultivated for agriculture. More recently, however, there has been a growing interest in restoring freshwater wetlands to prevent further soil subsidence. In one of the approaches used, the restored wetlands in the Delta are flooded with a water table that is above ground level at all times (Hemes et al., 2019). The four restored wetlands in the Delta selected for this study represent a range of conditions with some sites dominated by open water areas and others covered in dense vegetation throughout (Eichelmann et al., 2018), representing varying amounts of T/ET ratios expected at the different sites.

While restoring freshwater wetlands in the Delta can have many benefits, including those related to wildlife habitat, climate, recreation, and levee stability, it can also lead to increased water loss through ET depending on the vegetation cover characteristics (Eichelmann et al., 2018). Moreover, given that changes in local and regional ET can affect cloud formation and precipitation distribution (Gerken et al., 2018), this may have a knock-on effect on the water cycle and on the climate feedback of wetlands (Hemes et al., 2018). In locations that experience spatial and temporal water shortages, such as California, increasing our knowledge of the local water cycle and understanding how ET is affected by external drivers is extremely important.

Here, we show that we can partition ET measurements above flooded wetlands in the Delta by predicting daytime E from nighttime ET measurements using ANNs in combination with environmental driver variables such as vapor pressure deficit (VPD), temperature, atmospheric turbulence, canopy greenness index, and others. The meso-network of diverse wetland EC sites used in this study is ideal to test this new ET partitioning method as it provides a continuum of T/ET conditions across complex canopy architectures. We present the most promising models and discuss the application of ANN to partition T/ET measurements. While there is an emphasis on wetlands, we show evidence that our method may be applied to other ecosystems as well, increasing the knowledge of the water cycle and shedding light on plant-water productivity relationships at an ecosystem level.

## 2 Methods

### 2.1 Site Description

We conducted EC measurements at four wetland sites in the Sacramento-San Joaquin river delta in Northern California: West Pond (38° 6.44′N, 121° 38.81′W, Ameriflux ID: US-TW1), East End (38° 6.17′N, 121° 38.48′W, Ameriflux ID: US-TW4), Mayberry Farms (38° 2.99′N, 121° 45.90′W, Ameriflux ID: US-MYB), and Sherman Island (38° 2.21′N 121° 45.28′W, Ameriflux ID: US-Sne). All sites are part of the Ameriflux network and the EC data from these sites are available for download through the Ameriflux data sharing platform (https://ameriflux.lbl.gov/). The sites have been described in detail in other publications (Detto et al., 2010; Hatala et al., 2012; Knox et al., 2015; Eichelmann et al., 2018; Hemes et al., 2018, 2019) and their main characteristics will only be briefly summarized here. All four wetlands are artificially constructed wetlands managed by the Department of Water Resources to reverse soil subsidence in the area. The water table is actively managed to be above ground level throughout the flooded portions of the wetlands at all sites.

The West Pond wetland is the oldest of the four wetlands, originally constructed in 1998. It is the most homogeneous of the study sites, with a fairly even, but slightly sloping, ground surface and dense vegetation covering the whole wetland (97% vegetation cover within EC footprint in 2018, Valach et al., 2021). The water table varies slightly throughout the wetland due to the sloping ground level but is generally between 20 and 40 cm above ground level. The Mayberry Farms wetland was constructed in 2010 and has a very heterogeneous footprint. With a heterogeneous bathymetry this wetland features small islands of vegetation and deeper channels and pools of open water (64% vegetation cover within EC footprint in 2018, Valach et al., 2021). The water depth varies from 2 m above ground level to 2 cm above ground level in the flooded portions, with some dry areas. The East End wetland was constructed in 2013 and also features some areas of open water channels and pools. The vegetation at East End has filled in more evenly since its establishment and it has a greater vegetation cover than Mayberry Farms (96% vegetation cover within EC footprint in 2018, Valach et al., 2021). The Sherman Island wetland is the newest wetland constructed in 2016. Similarly to Mayberry Farms, it features a very heterogeneous bathymetry and the footprint is dominated by large portions of open water. Vegetation has only taken hold in very few and small patches within the footprint of the EC measurements (45% vegetation cover within EC footprint in 2018, Valach et al., 2021). While the individual make-up and proportions vary slightly between sites, the dominant vegetation species at all sites are tules (*Schoenoplectus acutus*) and cattails (*Typha* spp.) (O’Connell et al., 2015).

### 2.2 Eddy Covariance Data

We measured continuous fluxes of H_2_O, CO_2_ and sensible heat using the EC method at all sites (Baldocchi et al., 1988). A detailed description of the instrument set-up and calculation procedures can be found in previously published papers (Detto et al., 2010; Hatala et al., 2012; Knox et al., 2015; Eichelmann et al., 2018; Hemes et al., 2018, 2019) and will only be summarized here. At each site, the EC instrumentation consisted of a sonic anemometer (WindMaster 1590 or WindMaster Pro 1352, Gill Instruments Ltd, Lymington, Hampshire, England) and an open path trace gas analyzer for H_2_O and CO_2_ concentrations (LI-7500 or LI-7500A, LI-COR Inc., Lincoln, NE, USA). The instruments were mounted at a fixed height at least 1 m above the maximum height of the canopy.

High frequency (20 Hz) measurements of sonic temperature, three-dimensional wind speed, and trace gas concentrations were recorded on USB drives in the field through the analyzer interface (LI-7550, LI-COR Inc., Lincoln, NE, USA). The data were collected approximately every two weeks, with routine maintenance and servicing of the instruments taking place at the same time. The LI-7500 trace gas analyzers were calibrated approximately every three to six months in the laboratory. The performance of the EC set-up was also cross checked periodically at individual sites by the Ameriflux mobile EC reference system (Schmidt et al., 2012).

All data processing and filtering was performed offline. Thirty-minute average fluxes were calculated using custom software written in-house (MATLAB, MathWorks Inc., R2015b, version 8.6.0) after basic de-spiking of high frequency data and filtering for instrument malfunctioning (Detto et al., 2010; Hatala et al., 2012; Knox et al., 2015; Eichelmann et al., 2018). A rotation into the mean wind was performed for each 30-minute averaging interval and the Webb-Pearman-Leuning correction for air density fluctuations for open path sensors was applied to the calculated fluxes (Webb et al., 1980). Fluxes were filtered for low friction velocity (u_∗_), as well as based on stability and turbulence conditions (Foken & Wichura, 1996). Low friction velocity thresholds are based on the point where nighttime CO_2_ fluxes become independent of u_∗_and are defined individually at each site. The thresholds can vary seasonally and usually range from 0.12 m s^-1^ to 0.2 m s^-1^. Because of the narrow shape of the wetland, the West Pond wetland fluxes were also filtered by wind direction to ensure flux footprints originated from the ecosystem of interest.

Energy budget closure is often used as a quality indicator for EC data (Wilson et al., 2002). At the flooded wetland sites covered in this study the energy budget closure of daily totals was between 73% and 81%, which is slightly lower than typically found in dry ecosystems. H_2_O fluxes from the West Pond, Mayberry Farms, and East End wetland sites used in this study have been published and discussed in detail by Eichelmann et al. (2018), including a discussion of data quality, energy budget closure, and the difficulties estimating energy storage components in the flooded wetlands. Because of the importance of storage terms in the context of these sites, energy fluxes measured by the EC method have not been adjusted for incomplete energy budget closure (Eichelmann et al., 2018). In this study, positive fluxes indicate a gain to the atmosphere and negative fluxes indicate a loss from the atmosphere. All analyzes and data processing described in this study were performed using MATLAB (MathWorks Inc., R2018a, version 9.4.0).

### 2.3 Auxiliary Data

Meteorological and environmental data were also measured continuously in addition to EC data at all sites. The following auxiliary measurements were available at all wetland sites: Air temperature (T_air_); water temperature at 3 to 6 different water depths (T_water_, depths vary between site due to differences in water tables); soil temperature at 6 different depths (T_soil_); relative humidity (RH); atmospheric pressure; incoming and outgoing shortwave radiation; incoming and outgoing longwave radiation; net radiation; incoming and outgoing photosynthetically active radiation; water table depth; water conductivity; and vegetation greenness index from camera data (GCC). Moreover, the West Pond and East End wetland sites were equipped with a rain gauge to measure precipitation and the East End wetland site was equipped to measure ground heat flux (G).

Data were recorded as half hour averages (or totals in the case of precipitation) with individual sampling frequency varying between 1 and 15 minutes depending on the sensor. Specifically of interest for this study are measurements of vapor pressure deficit (VPD), water table depth (WT), air temperature (T_air_), vegetation greenness index (green chromatic coordinate; GCC), and net radiation (R_net_). VPD was calculated from relative humidity measurements in combination with air temperature data, both measured with aspirated and wind-shielded humidity and temperature probes (HMP-60, Vaisala Inc., Helsinki, Finland). Net radiation was measured using either a net radiometer (NR-LITE Radiometer, Hukseflux, Delft, the Netherlands; at Mayberry Farms) or a four-component net radiometer (NR01 Net Radiometer, Hukseflux, Delft, the Netherlands; at West Pond, East End, and Sherman Island).

### 2.4 Artificial Neural Network Partitioning Routine

Artificial Neural Networks have been applied for gap-filling and partitioning EC fluxes in the past (Papale & Valentini, 2003; Oikawa et al., 2017; Tramontana et al., 2020). Specifically, for CO_2_ fluxes, ANNs have shown to perform well when used to gap-fill missing data (Moffat et al., 2007) and partitioning net CO_2_ fluxes into the component fluxes of gross primary production (GPP) and ecosystem respiration (R_eco_) (Desai et al., 2008; Oikawa et al., 2017; Tramontana et al., 2020). Following a similar approach to partitioning CO_2_ data, we assumed that nighttime ET data is dominated by E at these flooded sites:

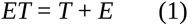

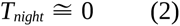

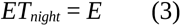

We conducted several leaf-level chamber measurements using a LI-6400 Portable Photosynthesis System (LI-COR Inc., Lincoln, NE, USA) throughout the growing season of 2017 to confirm that nighttime and dark T flux is indeed negligible at these sites. The available nighttime E data is used in combination with environmental input variables to train the ANN routine to predict daytime E. Daytime T was then calculated as the difference between total ET and E:

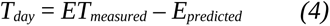

Before ET partitioning was performed all flux data were gap-filled using ANN routines described in previous studies (Knox et al., 2015, 2016; Oikawa et al., 2017, Eichelmann et al., 2018).

#### 2.4.1 Artificial Neural Network Routine Set-up

To partition ET data using ANNs in this study, we followed a similar set-up and architecture as described for gap-filling and partitioning CO_2_ data in previous studies (Baldocchi & Sturtevant, 2015; Knox et al., 2015, 2016; Oikawa et al., 2017). The entire available (multi-year) explanatory dataset was split into 20 data clusters using the k-means clustering algorithm. The data used for training, testing, and validation of the ANNs was proportionally sampled from these clusters with one third of the available data used for training, testing, and validation each. This procedure avoids a sampling bias towards periods when more data are available, such as a specific time of the year or time of the day. Proportional data sampling from the k-means clusters into training, testing, and validation data was repeated 20 times. For each of the 20 re-sampled training, testing, and validation datasets several ANN architectures were tested starting with one hidden layer and the same number of nodes as the number of explanatory input variables (n_inputvar_). Each architecture was initialized 10 times with random starting weights and the initialization with the lowest mean sampling error was used. The complexity of the ANN architecture was increased first by increasing the number of nodes to 1.5 times n_inputvar_ and then by increasing the number of hidden layers until a further increase in complexity results in less than 5% reduction of the mean standard error. For our datasets, this commonly resulted in the use of an architecture with two hidden layers, the first one with n_inputvar_ nodes, the second one with 0.5*n_inputvar_ nodes, although for some sites and input variable combinations architectures with only one hidden layer produced better results. The ‘validation’ step within the ANN procedure described above is performed on nighttime data only and is therefore distinctly different from the validation with flooding and leaf level data described below. Throughout the remainder of the manuscript when we use the term ‘validation’ we refer to the independent flooding and leaf level data validation. The ANN internal validation routine based on nighttime data is referred to as ‘testing’.

#### 2.4.2 Selection of Explanatory Variables

A number of different explanatory environmental input variables were tested individually and in combination. Based on the general understanding of the drivers of E fluxes in terrestrial and aquatic ecosystems we tested the following input parameters: Meteorological and environmental variables: VPD, R_net_, GCC, WT, T_air_; Flux variables: friction velocity (u_∗_), gap-filled sensible heat flux (H_gf), gap-filled CO_2_ flux (wc_gf), and ecosystem respiration (er_Reichstein) partitioned using the temperature dependency method proposed by Reichstein et al. (2005). In addition, we used a running decimal timestamp (datetime) as input variable in all our ANN runs. VPD, u_∗_, and T_air_ describe the atmospheric demand driving E. Rnet and H_gf are connected to ET (or latent energy) through the energy balance equation. GCC, wc_gf, and er_Reichstein are directly or indirectly related to plant physiological responses that can impact ET components. Finally, WT is related to the water budget of the ecosystem. Given the strong correlation of water temperature (T_water_) with nighttime ET documented at these sites in a previous study (Eichelmann et al., 2018) we would also expect T_water_ to perform well as an environmental input variable. Unfortunately, we were unable to include T_water_ as an input variable in this study since we did not have consistent T_water_ measurements across time for any of the four sites.

We ran the ANN routine for each of these parameters individually and recorded the R^2^ value and slope of the linear regression of the nighttime EC data initially set aside for testing within the ANN routine versus the predictions. This R^2^ value is called ‘testing R^2^’ throughout this manuscript and is based only on nighttime data. Starting with the input parameter with the highest testing R^2^, we ran the ANN routine with increasing numbers of input variables, each time adding on the variable with the next highest testing R^2^ value. We continued this process until a further increase in input variables resulted in less than 1% increase in the testing R^2^ value. We averaged the testing R^2^ values across the four sites and used this value to estimate increases in the performance of the ANNs. While this average testing R^2^ does not have any statistical relevance, it gave us a good indicator on how well the models performed across all sites studied.

### 2.5 Validation of Results

One of the main issues facing validation of ET partitioning methods is often the lack of independent E or T data to validate against (Stoy et al., 2019). Taking independent measurements of ecosystem E or T is challenging and one of the main reasons why partitioning approaches for EC measurements of ET are much sought after. Since we do not have independent measurements of ecosystem level E or T available at our sites, we reverted to validating our partitioning data by a conditional sampling approach, selecting EC measurement data from certain time periods when E and T can be known or closely approximated to compare with the ANN predicted E or T. One of these time periods is the initial time right after flooding of the wetland (referred to as flooding data), when vegetation had not yet established within the footprint of our instruments. During this time, it can be assumed that the entire H_2_O flux coming off the surface is from E, with negligible T.

Since we trained our ANN routines only on nighttime data, we were able to use the daytime data during the initial flooding period as an independent validation dataset for E. Apart from the initial flooding period, T can also be assumed to be small to negligible during the senescent winter months. However, since the plants are not harvested or otherwise removed and the climate in this region is fairly mild, some do stay green throughout the winter and may continue to be photosynthetically active. Additionally, vegetation on dry areas such as levees usually starts to green up during the winter months in this region. Both of these would be contributing to a small T flux from the ecosystem. Moreover, ET fluxes during the winter period are generally lower and subject to larger errors due to more challenging turbulence conditions during this time. Such conditions result in large relative error in flux measurements during this period limiting the insights gained from the validation during the senescent winter period. Nonetheless, we included validation of E predicted from our ANN method against E measured during winter times to further test the performance of our method.

In addition to the validation during periods when T was zero, we also conducted a number of leaf-level T measurements in the summer of 2017 at the East End wetland using a LI-6400 portable photosynthesis system (LI-COR Inc., Lincoln, NE, USA) with a clear conifer chamber (part number 6400-05) encasing sections of the leafs or culms. Six individual leaf-level measurement points (three for each of the dominant plant species) taken during the same half hour period were pooled to allow comparison with the half hourly EC data. These measurements provided us with an estimate of T per unit of sunlit leaf area and may potentially be converted to the ecosystem scale if the ecosystem leaf area index and the leaf angle distribution are known. Efforts have been made to estimate the leaf area index in a number of the wetlands in the study region, however, due to the high heterogeneity and litter accumulation in these systems there is a high level of uncertainty associated with the measured leaf area indexes (Dronova & Taddeo, 2016). Additionally, the leaf angle distribution is unknown in these systems and can only be approximated, which is an intrinsic limitation of this technique.

Taking all these uncertainties into account, ecosystem T scaled up from leaf-level measurements is associated with very large error intervals and cannot serve as a reasonable constraint on the absolute values of our ANN partitioned T fluxes. However, since the scaling factors to convert leaf-level values to ecosystem level are constant multipliers, we should still be seeing a linear relationship between the leaf-level flux and the partitioned ecosystem level T if our partitioning algorithm predicts the correct T behavior across a range of environmental conditions. While we may not be able to compare the absolute T values, we can compare the response cycle of ANN predicted T with the field measurements to validate that we are predicting the right behavior.

### 2.6 Comparison with Other T/ET Partitioning Approaches

Direct comparisons with the Scott and Biederman’s (2017) method were carried out in order to evaluate the performance of our own models against their approach. For that, we used the model (F11, see Results below) that achieved the best R^2^ value against the validation with leaf-level/flooding data. While Scott and Biederman (2017) forced all monthly regressions between ET and gross ecosystem productivity (GEP) to the same slope, we used different slopes for each regression. This was done to ensure the best fitting since our datasets did not show the same uniform behavior across months. Indirect comparisons with other methodologies mentioned above were also discussed.

## 3 Results

### 3.1 Artificial Neural Network Architecture Performances

Alongside the basic timestamp (datetime), VPD and T_air_ were the meteorological variables that best explained our data when only looking at the nighttime testing data, with average testing R^2^ values across all sites of 0.648 and 0.565, respectively (Table 1 and Supplementary Table 1). The flux related variables that showed the highest average testing R^2^ values and added most information to the models were H_gf (testing R^2^ of 0.620) and u_∗_ (testing R^2^ of 0.531). To increase the ANNs complexity we, therefore, followed the variables order of VPD > H_gf > T_air_ > u_∗_, adding each of them into the models sequentially. VPD was the variable that contributed the most to increase the testing R^2^ values of the ANNs, with an average increase of 24% across all sites and a maximum of 36% for West Pond, when models F21 and F26 were compared (Table 1). The incorporation of H_gf was responsible for an average increase of 10% in testing R^2^, when comparing the ANNs F26 and F33 (Table 1). T_air_ only increased the ANNs testing R^2^ by 1% (i.e., when comparing models F33 and F34), however, when we added u_∗_, the average testing R^2^ value increased across all sites by 9%, when comparing models F34 and F11 (Table 1). Thus, building the ANN F11 using datetime, VPD, H_gf, T_air_, and u_∗_, the average testing R^2^ value across all sites reached 0.853, with a minimum of 0.728 (West Pond) and a maximum of 0.910 (Sherman Island; Supplementary Table 1).

**Table 1:** Average testing R^2^ and slope values for 12 ANN architecture models used to partition evapotranspiration measurements, demonstrating an increase in complexity from models F21 (most basic) to F36 (most complex).

| Model Name | Model Structure | Average testing $R^2$ | Average Slope |
| --- | --- | --- | --- |
| F21 | datetime | 0.410 | 0.393 |
| F26 | datetime + VPD | 0.648 | 0.626 |
| F17 | datetime + VPD + $T_{air}$ | 0.672 | 0.636 |
| F31 | datetime + VPD + $T_{air}$ + GCC | 0.686 | 0.657 |
| F32 | datetime + VPD + $T_{air}$ + GCC + Rnet | 0.689 | 0.665 |
| F15 | datetime + VPD + $T_{air}$ + GCC + Rnet + WT | 0.694 | 0.663 |
| F33 | datetime + $H_{gf}$ + VPD | 0.753 | 0.726 |
| F34 | datetime + $H_{gf}$ + VPD + $T_{air}$ | 0.762 | 0.734 |
| F11 | datetime + $H_{gf}$ + VPD + $T_{air}$ + $u_*$ | 0.853 | 0.831 |
| F35 | datetime + $H_{gf}$ + VPD + $T_{air}$ + $u_*$ +<br>er_Reichstein | 0.863 | 0.851 |
| F14 | datetime + $H_{gf}$ + $u_*$ + VPD + $T_{air}$ + GCC +<br>Rnet + WT | 0.877 | 0.868 |
| F36 | datetime + $H_{gf}$ + $u_*$ + wc_gf + er_Reichstein +<br>VPD + $T_{air}$ + GCC + Rnet + WT | 0.891 | 0.880 |

Of all the 36 ANNs tested, the highest average testing R^2^ (0.891) was reached when all the explanatory variables (i.e., datetime, H_gf, u_∗_, wc_gf, er_Reichstein, VPD, T_air_, GCC, Rnet and WT) were put into the model F36 (Table 1 and Supplementary Table 1). Consequently, on average, all the other variables analyzed (i.e., wc_gf, er_Reichstein, GCC, Rnet and WT) accounted for less than 4% of the testing R^2^ value across all the four sites (when comparing models F36 and F11; Table 1). The top five ANNs (F36 > F14 > F20 > F35 > F11) that performed better than 0.85 all have datetime, VPD, H_gf, T_air_, and u_∗_as their explanatory variables and all the 11 ANNs that scored an average testing R^2^ higher than 0.80 have both VPD and u_∗_in their models (Table 1 and Supplementary Table 1). Fifteen ANNs showed an average testing R^2^ higher than 0.70 and the lowest average testing R^2^ among these (0.730) was presented by the ANN F2, constructed using only datetime, T_air_, and u_∗_ (Supplementary Table 1). Unsurprisingly, the lowest average testing R^2^ (0.410) of all the 36 ANNs analyzed was given by the ANN built using datetime alone (F21). The slope values (Table 1 and Supplementary Table 2) of the different ANNs followed quite closely the pattern described for the increase in testing R^2^ values.

### 3.2 Validation of Artificial Neural Networks

#### 3.2.1 Flooding Validation

To evaluate the performance of our ANN partitioning method, we compared the model predicted E with EC measurement data from conditionally sampled post-flooding periods, during which we assume T to be negligible (Table 2). The ANN F11 showed the highest validation R^2^ values for East End (0.81), Mayberry Farms (0.69), and Sherman Island (0.82). These values surpassed those from the model F36 (most complex), which reached 0.51, 0.56, and 0.53, for East End, Mayberry Farms, and Sherman Island, respectively. Figure 1 shows the validation comparison between F11 and F36 for the three sites.

**Figure 1:**
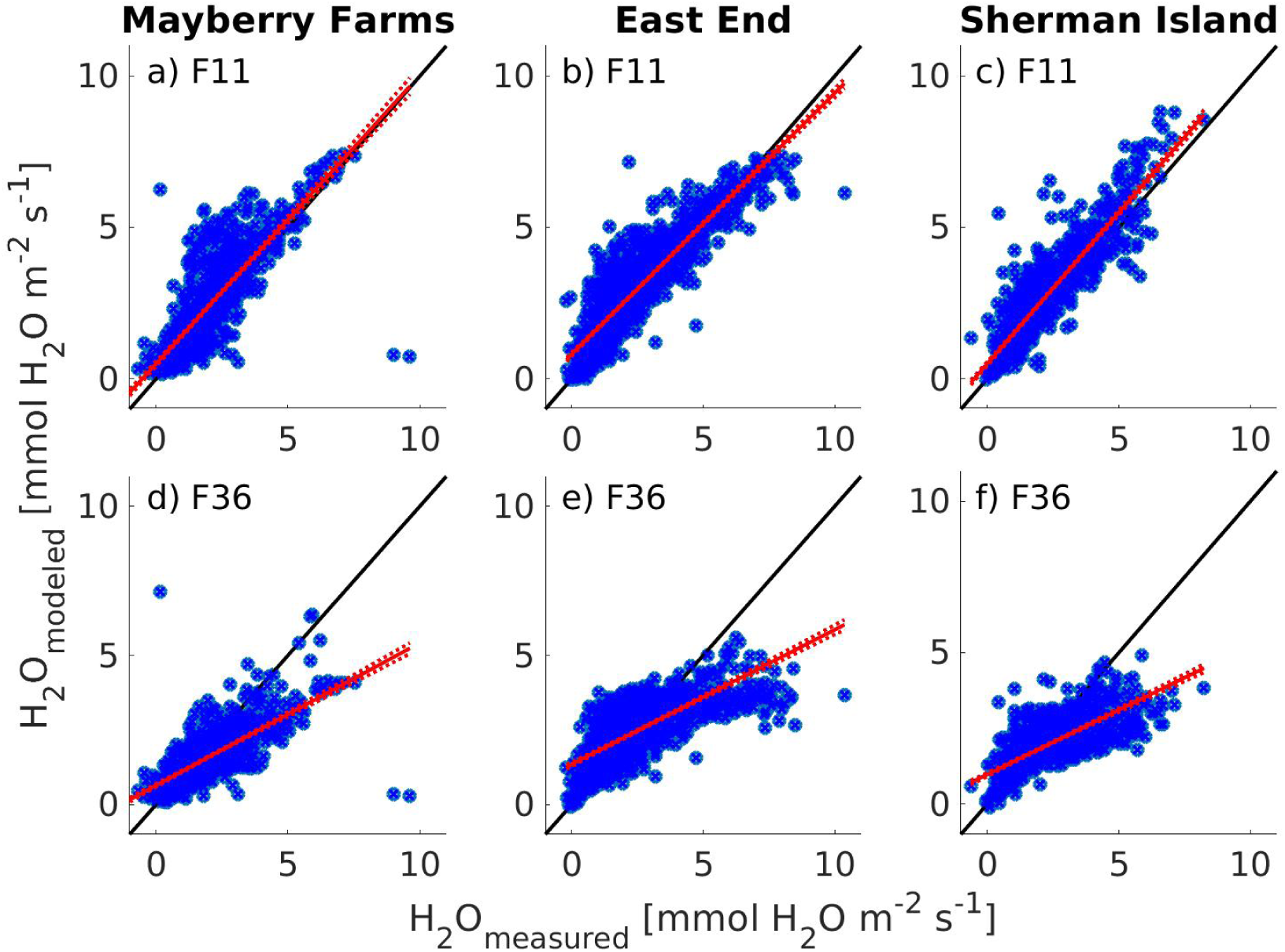
Comparison between the eddy covariance measured daytime evaporation flux (H_2_O_measured_) and daytime evaporation predicted by ANNs (H_2_O_modeled_) using model F11 (top panels, a-c) and F36 (bottom panels, d-f) based on data collected right after flooding for Mayberry Farms (a, d), East End (b, e), and Sherman Island (c, f). Note: the black lines are 1:1 relationships for reference, red lines show linear regressions with standard deviation, and blue dots represent the data.

**Table 2:** Validation R^2^ and slope values of seven ANNs used to partition evapotranspiration measurements and validated with data collected right after flooding for East End, Mayberry Farms, and Sherman Island wetland sites. Models are ordered by the increase in complexity, from model F21 (most basic) to F36 (most complex). Refer to Tables 1 and 3 for each model’s input variables. Validation R^2^ values higher than 0.7 are highlighted in bold.

| Model Name | East End |  | Mayberry Farms |  | Sherman Island |  |
| --- | --- | --- | --- | --- | --- | --- |
| | $R^2$ | Slope | $R^2$ | Slope | $R^2$ | Slope |
| F21 | 0.29 | 0.28 | 0.06 | 0.09 | 0.34 | 0.25 |
| F26 | 0.48 | 0.52 | 0.26 | 0.37 | 0.61 | 0.50 |
| F17 | 0.50 | 0.46 | 0.31 | 0.41 | 0.63 | 0.56 |
| F15 | 0.24 | 0.15 | 0.16 | 0.13 | 0.37 | 0.28 |
| F33 | 0.61 | 0.66 | 0.48 | 0.81 | 0.62 | 0.71 |
| F11 | **0.81** | 0.86 | 0.69 | 0.95 | **0.82** | 1.00 |
| F36 | 0.51 | 0.45 | 0.56 | 0.48 | 0.53 | 0.43 |

#### 3.3.2 Winter Time Validation

Judging by the observed R^2^ values, the validation using daytime data from senescent periods during the winter time (December to February, Table 3) performed quite poorly in comparison to the validation performed with data during the initial flooding periods (Table 2). Nevertheless, the winter period validation overall did confirm the same trends and observations as the flooding validation. At Mayberry Farms and Sherman Island ANN F11 again had the highest R^2^ values (0.56 and 0.70, respectively). However, at East End and West Pond the model F36, which included all input variables, performed best with R^2^ values of 0.45 and 0.36, respectively. Figure 2 shows the validation comparison between F11 and F36 for the four sites using winter data.

**Figure 2:**
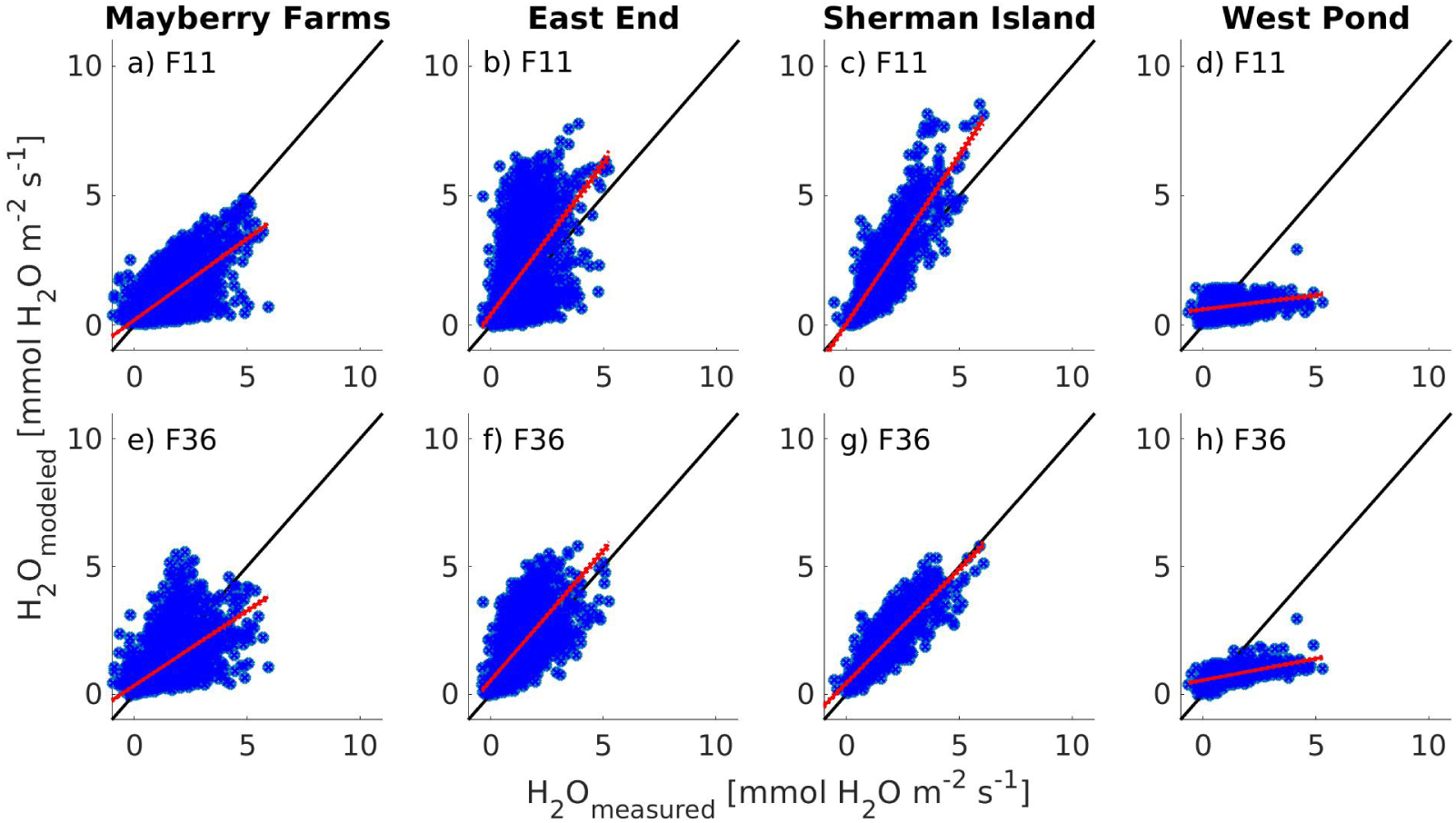
Comparison between the eddy covariance measured daytime evaporation flux (H_2_O_measured_) and daytime evaporation predicted by ANNs (H_2_O_modeled_) using model F11 (top panels, a-d) and F36 (bottom panels, e-h) based on data collected during senescent periods in winter (December to February) at Mayberry Farms (a, e), East End (b, f), Sherman Island (c, g), and West Pond (d, h). Note: the black lines are 1:1 relationships for reference, red lines show linear regressions with standard deviation, and blue dots represent the data.

**Table 3:** Validation R^2^ and slope values of seven ANNs used to partition evapotranspiration measurements and validated with winter time data (December to February) for each of the four wetlands studied (East End, Mayberry Farms, Sherman Island, and West Pond). Models are listed according to the increase in complexity, from model F21 (most basic) to F36 (most complex). Refer to Tables 1 and 4 for each model’s input variables. Validation R^2^ values higher than 0.7 are highlighted in bold.

| Model<br>Name | East End |  | Mayberry Farms |  | Sherman Island |  | West Pond |  |
| --- | --- | --- | --- | --- | --- | --- | --- | --- |
| | $R^2$ | Slope | $R^2$ | Slope | $R^2$ | Slope | $R^2$ | Slope |
| F21 | 0.06 | 0.02 | 0.06 | 0.06 | 0.15 | 0.11 | 0.08 | 0.30 |
| F26 | 0.17 | 0.25 | 0.26 | 0.38 | 0.45 | 0.48 | 0.03 | 0.08 |
| F17 | 0.21 | 0.24 | 0.35 | 0.47 | 0.47 | 0.49 | 0.05 | 0.06 |
| F15 | 0.33 | 0.41 | 0.14 | 0.12 | 0.43 | 0.29 | 0.17 | 0.11 |
| F33 | 0.21 | 0.71 | 0.22 | 0.48 | 0.19 | 0.42 | 0.01 | 0.01 |
| F11 | 0.33 | 1.15 | 0.56 | 0.71 | **0.70** | 1.27 | 0.11 | 0.11 |
| F36 | 0.45 | 0.95 | 0.43 | 0.59 | 0.69 | 0.87 | 0.36 | 0.17 |

#### 3.3.2 Validation on Diurnal Measurements of Leaf-Level Data for East End

To evaluate the performance of our method further, we compared the model predicted T with independent leaf-level data collected during a field campaign in summer 2017 at the East End wetland. The leaf-level data showed high variability across individual measurements (Fig. 3). F11 again showed a high R^2^ (0.986, Table 4). Other models (F15, F33) also performed quite well in the leaf-level validation, in contrast to their performance for the validation during flooding or senescent periods. The most complex ANN (F36) had a lower R^2^ value (0.92) for the leaf-level validation. In general, adding too many variables did not lead to enhancement of validation values, but it is to be noted that all models showed a high level of agreement with the leaf-level data (Table 4). Figure 3 shows both F11 and F36 validations against leaf-level data.

**Figure 3:**
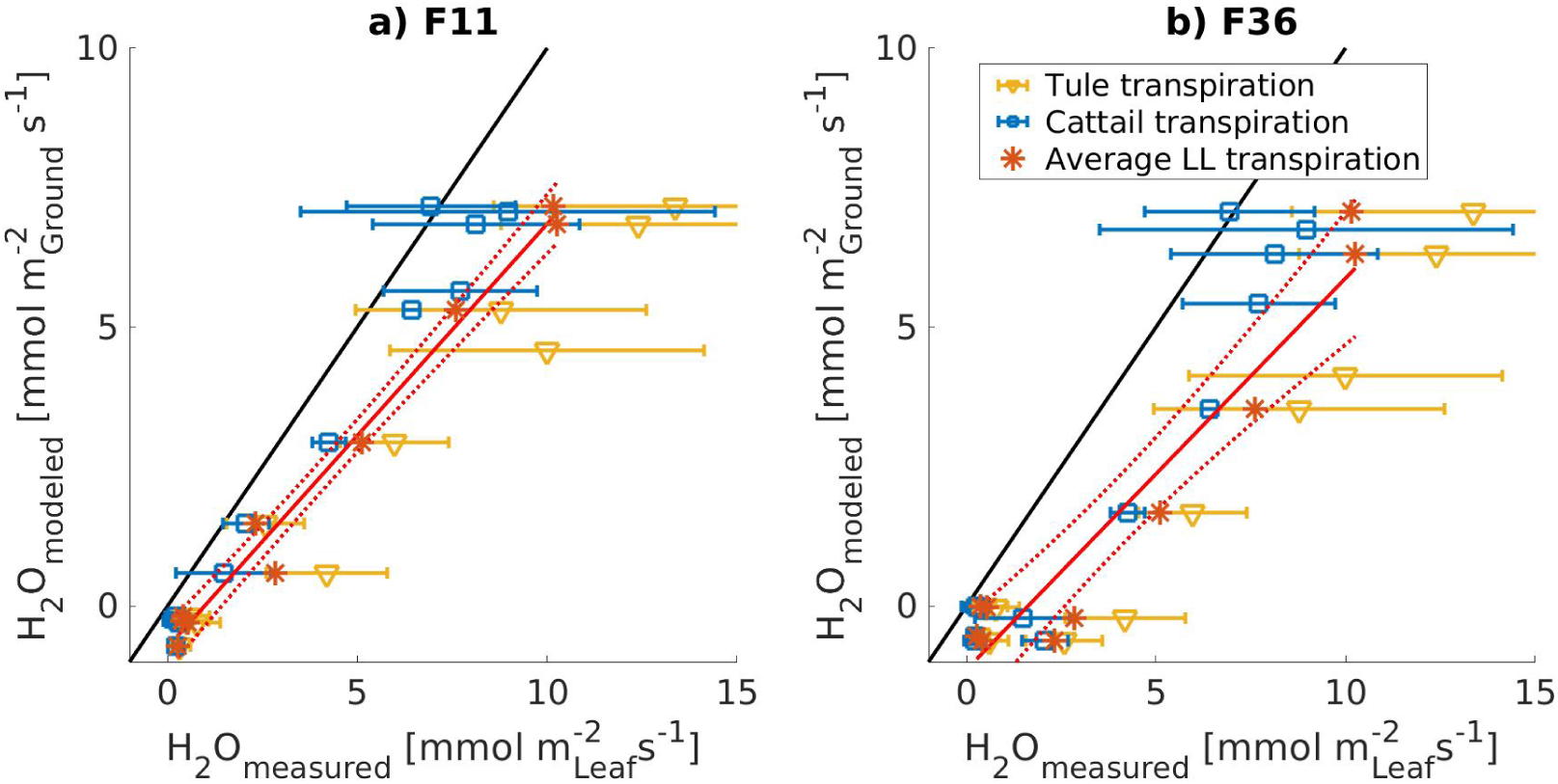
Ecosystem level transpiration data (H_2_O_modeled_) predicted by ANNs F11 (a) and F36 (b) validated against leaf-level transpiration data (H_2_O_measured_) collected during the field campaigns in 2017 for the two dominant species in the wetland: Tule (yellow triangles) and Cattail (blue squares). The overall linear regression line (solid red line) and standard deviation (dashed red line) is based on average leaf-level transpiration across both species (red asterisks). Error bars represent the standard deviation from the mean for each measurement interval and species for the leaf-level data. Leaf-level data were pooled for 30 min intervals to match the eddy covariance averaging period. The solid black lines show 1:1 relationships for reference.

**Table 4:** R^2^ and slope values for linear regression of ecosystem level transpiration data predicted by seven ANNs versus leaf-level transpiration data collected in 2017 for East End. Models are ordered by the increase in complexity from model F21 (most basic) to F36 (most complex).

| Model Name | Model Structure | $R^2$ value | Slope value |
| --- | --- | --- | --- |
| F21 | datetime | 0.979 | 0.95 |
| F26 | datetime + VPD | 0.984 | 0.79 |
| F17 | datetime + VPD + TA | 0.984 | 0.75 |
| F15 | datetime + VPD + TA + GCC + Rnet + WT | 0.987 | 0.81 |
| F33 | datetime + H_gf + VPD | 0.99 | 0.93 |
| F11 | datetime + H_gf + VPD + TA + $u_*$ | 0.986 | 0.76 |
| F36 | datetime + H_gf + $u_*$ + wc_gf + er_Reichstein +<br>VPD + TA + GCC + Rnet + WT | 0.922 | 0.70 |

### 3.3 Artificial Neural Networks Performance Across the Wetland Sites

To look for model consistency across diverse canopy architecture and successional stages, we compared ANN testing R^2^ values between the four sites. Among the four sites, East End and Sherman Island were the only sites that had ANNs with testing R^2^ values larger than 0.90 for the EC testing data set aside during the ANN routine (Supplementary Table 1). At Sherman Island, East End, and Mayberry Farms 22, 20, and 19 ANN models reached testing R^2^ values above 0.70, respectively, whereas at West Pond only 11 models reached testing R^2^ values above 0.7 (Supplementary Table 1). In comparison with the other three studied sites, West Pond showed testing R^2^ values in the order of 9-18% smaller when analyzing the top five ANNs with average testing R^2^ larger than 0.85 (Supplementary Table 1). Considering all 36 ANNs, differences in testing R^2^ between the same ANN for different sites reached a maximum of 46%, when comparing model F6 at West Pond with Sherman Island (Supplementary Table 1).

### 3.4 Comparisons with Other Partitioning Approaches

To compare our ANN method with existing T/ET partitioning methods, we applied the Scott and Biederman (2017) long-term flux data partitioning method at all four sites. As expected, the Scott and Biederman (2017) method worked better for datasets with > 6 years (Fig. 4; Mayberry Farms, West Pond, and East End). Sherman Island, the shortest dataset with four years of data collection, performed poorly, showing negative correlations of ET vs GEP for the months of June to September (Fig. 4 d). Average monthly T fluxes from the Scott and Biederman (2017) method for Mayberry Farms and Sherman Island (Fig. 5a and d) both showed increases in T at the end of the growing season (i.e., October) out of line with the observed GEP patterns. Conversely, West Pond and East End (Fig. 5b and c) showed a T pattern parallel to GEP with the growing season.

**Figure 4:**
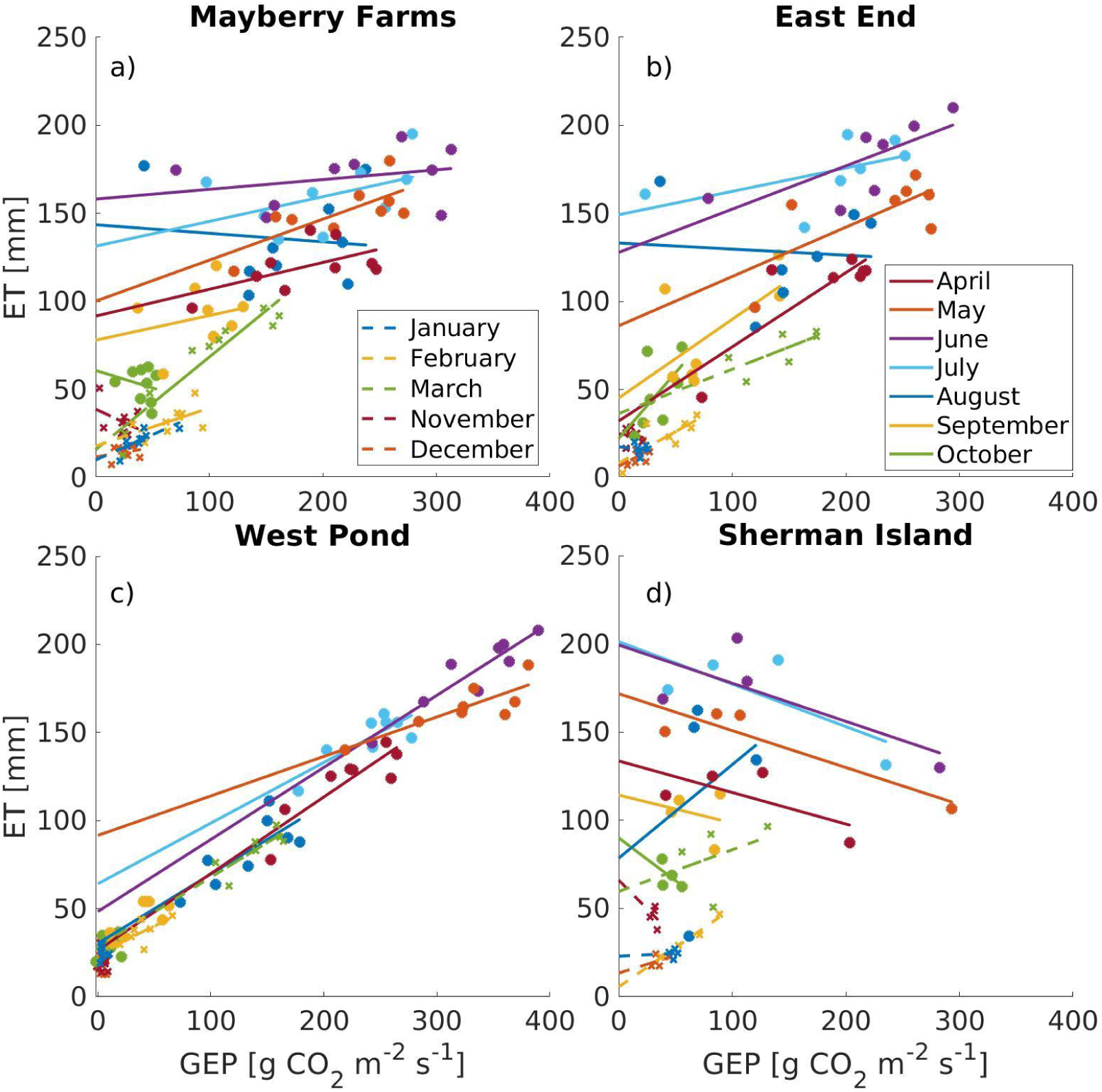
Monthly regressions of evapotranspiration (ET) vs Gross Ecosystem Productivity (GEP) data for four wetland sites Mayberry Farms (a), East End (b), West Pond (c), and Sherman Island (d) for T/ET partitioning using the Scott and Biederman (2017) method for long-term flux data. Each regression line represents data for the same month across multiple years. The method is considered unreliable for winter months when GEP is small (November through March, shown in dashed lines and cross symbols). Negative regression lines for most months at Sherman Island (d) indicate that the methodology does not work at this site, potentially due to the shorter time period of this dataset (4 years) or because of the large contribution of evaporation at this site (see main text for detailed discussion).

**Figure 5:**
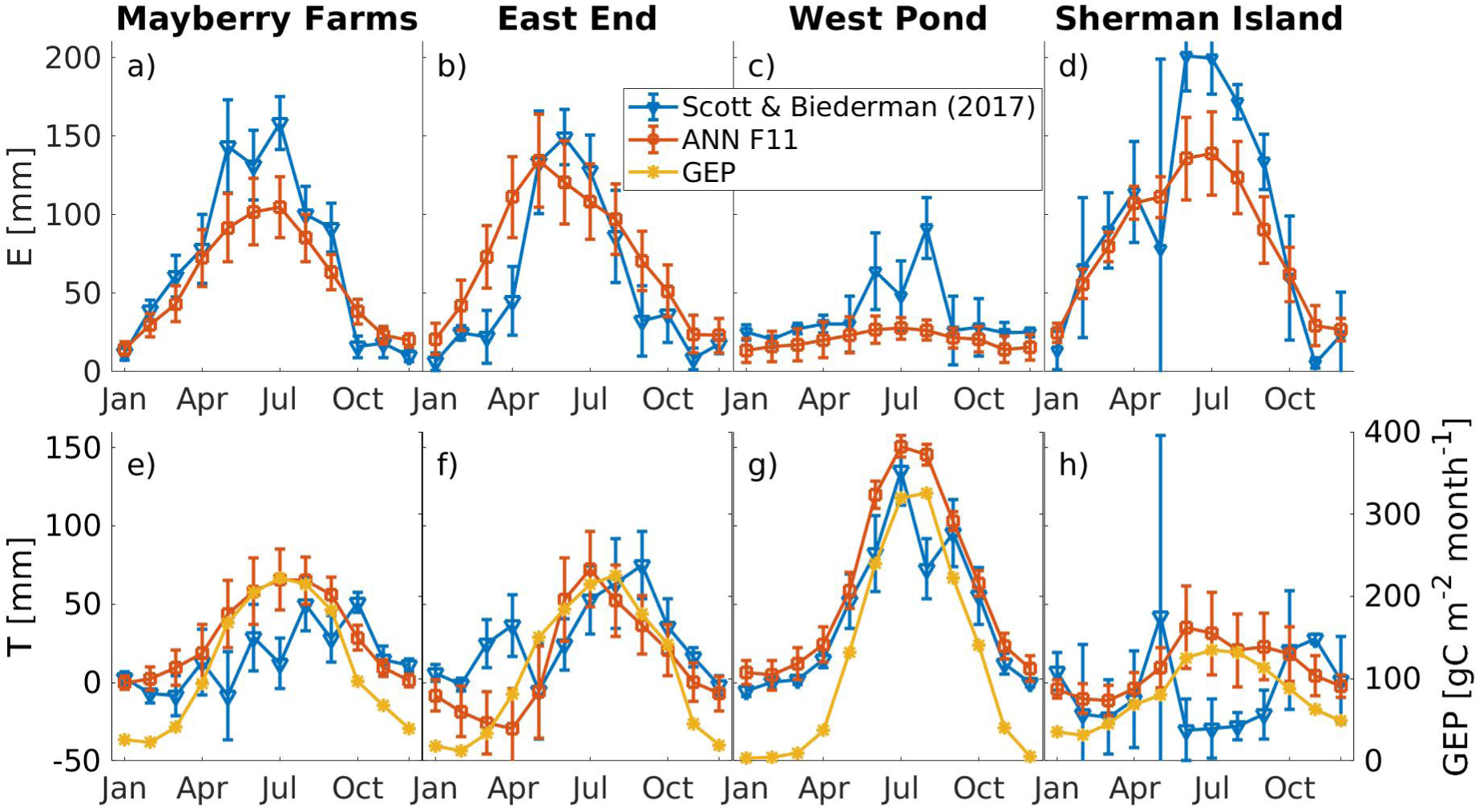
Average monthly evaporation (E) (top panels, a-d) and transpiration (T) (bottom panels, e-h) fluxes across four wetland sites: Mayberry Farms (a, e), East End (b, f), West Pond (c, g), and Sherman Island (d, h) comparing the ANN T/ET partitioning method described in this paper (red lines and square symbols) and the Scott and Biederman (2017) method (blue lines and triangle symbols) on long-term flux data. Error bars are based on the standard error of the fit intercept and slope for the Scott and Biederman (2017) method and on the interquartile range of the 20 individual ANN runs for the ANN method. Comparisons were done using ANN F11 for all sites. Gross Ecosystem Productivity (GEP, yellow lines and asterisks) for each site is shown for comparison in the bottom panels with a separate y-axis on the right.

While the T values from our ANN approach showed a similar behavior as GEP during the growing season, as would be expected, the T values from the Scott and Biederman (2017) method did deviate somewhat from the GEP pattern for all sites (Fig. 5). The best ANN (F11) also produced more reasonable T numbers for Sherman Island compared to the Scott and Bierderman (2017) method. In addition, the E values retrieved in our analysis for all sites were also more stable and did not fluctuate as much across months compared to the E values from the Scott and Biederman (2017) method (Fig. 5). While the Scott and Biederman (2017) method is not intended to produce reliable results for T/ET partitioning during winter months when GEP is small, it did show very good agreement of produced E and T values when compared to our ANN based values from October to February for all sites.

### 3.5 Resulting Evaporation and Transpiration Estimates

Figure 6 shows the annual (2013-2019) ANN based T/ET partitioning intercomparison for all sites using ANN F11. Only years with a full year of data are used. While ET stayed fairly consistent between 850-1250 mm for all sites and years (Fig. 6a), GEP showed more fluctuations between the different sites, as well as interannually within each site (Fig. 6b). Looking at the predicted partitioning of E and T (Fig. 6c, d), Sherman Island showed the highest values of E (approximately 1100 mm) for the three years of measurements available at this site, while West Pond had the lowest E values across all years and sites (200 to 300 mm). Although values at East End were always higher compared to Mayberry Farms for all years with measurements from both sites, decreasing pattern can be observed for E at both sites, ranging from high values of 831 mm at Mayberry Farms in 2013 and 1119 mm at East End in 2014 down to low values of 449 mm at Mayberry and 630 mm at East End in 2019. Transpiration showed opposite trends compared to E, with West Pond having the highest values (between 700-800 mm in most years), followed by Mayberry Farms with T values between 300-500 mm. The T pattern predicted at Mayberry Farms follows a similar pattern as the GEP measurements, most notably is the significant reduction in GEP in 2016 which was caused by saltwater intrusion at the site (Eichelmann et al., 2018, Chamberlain et al., 2020). This was mirrored in a reduction of T values in 2016, however, E was not affected. Sherman Island and East End showed T values below 300 mm for all years, considerably lower than the other two sites. In the first full year of measurements (2014), T at East End was even predicted as negative (−24 mm), similar to the negative T predictions observed at East End during the winter validation (Fig. 2). However, this value falls within the uncertainty range of 91 mm for annual ET measurements at this site in 2014 (Eichelmann et al., 2018). East End and Sherman Island both had a very high open water surface area, especially in the first years after flooding, so it would be expected that E is more dominant. Sherman Island specifically had extremely sparse vegetation cover throughout the EC measurement footprint for the first two years of measurements, also evident in the very low values of GEP. For both of these sites, East End and Sherman Island, we can see that gradually E declines and T increases as the vegetation fills in from year to year. Consequently, when comparing the T/ET values across sites (Fig. 6e), West Pond had the highest value of T/ET (70%-75% on T), followed by Mayberry Farms (30%-50%), East End (0-30%), and Sherman Island (<15%). This highlights that only West Pond can be described as a T dominated site with T/ET values in the range between 0.5 and 0.8 reported for other terrestrial ecosystems (Schlesinger & Jasechko, 2014). The other three sites are clearly E dominated and have T/ET values considerably lower than those expected for terrestrial ecosystems.

**Figure 6:**
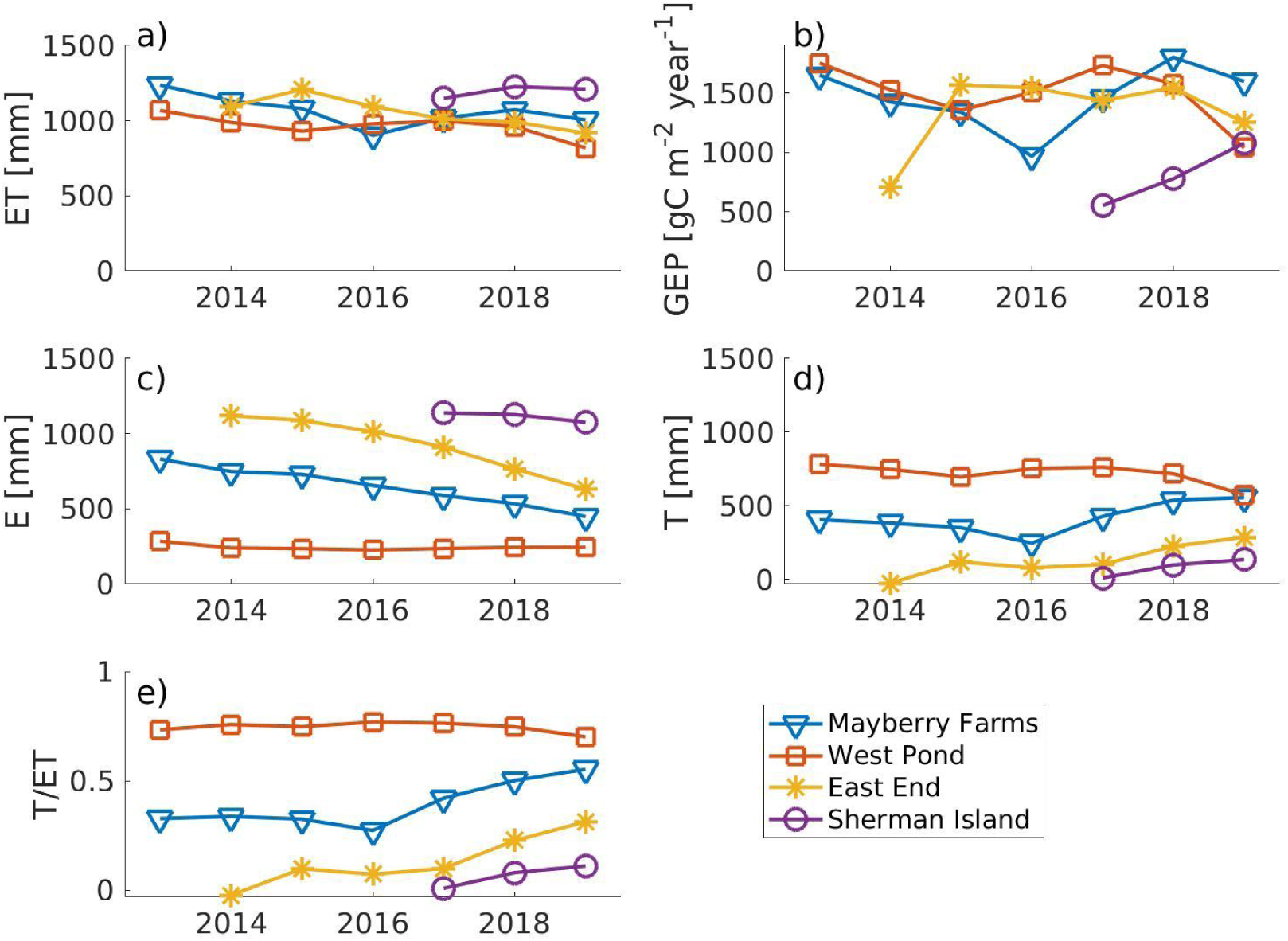
Annual intercomparison of (a) total evapotranspiration (ET), (b) gross ecosystem productivity (GEP), (c) evaporation (E), (d) transpiration (T), and (e) transpiration over evapotranspiration ratio (T/ET) between four wetland sites (Mayberry Farms, 2013-2019, blue triangles; West Pond, 2013-2019, red squares; East End, 2014-2019, yellow asterisks; and Sherman Island, 2016-2019, purple circles). E and T values are based on the ANN partitioning routine (F11) described in this study.

## 4 Discussion

### 4.1 Artificial Neural Network Architecture Performances

The ANN F36, which was built using all studied variables, presented the highest average testing R^2^ value (0.891) for the nighttime-based testing dataset among all 36 ANNs analyzed. Nevertheless, there was not much improvement in testing R^2^ in the models (i.e., maximum of 3-4% on average) after the ANN F11. This indicates that not all variables are necessary to provide good results in the partitioning of ET into E and T, and that less complex models can result in good predictions. For instance, using only datetime + H_gf + VPD (F33) or datetime + u_∗_+ T_air_ (F2) the average testing R^2^ value across all sites was > 0.70, indicating a good correlation. In addition, when using datetime + VPD alone the average testing R^2^ value for three sites (i.e., East End, Mayberry Farms and Sherman Island) was > 0.70.

In our study, the order of variable inclusion to increase model complexity was: datetime > VPD > H_gf > T_air_ > u_∗_. VPD was the variable that contributed the most in the improvement of the ANNs, with an average of 24% increase in testing R^2^ values across all sites. VPD is routinely measured at most EC sites (e.g., Fluxnet.org, 2021) and its effect on ecosystem water cycling by limiting surface conductance and reducing transpiration under high VPD is well documented (Buckley, 2005, Novick et al., 2016). The fact that the top 14 ANNs (i.e., with the highest testing R^2^ value) were constructed using VPD as one of the input parameters highlights the importance of VPD as a predictor of ecosystem water exchange. In addition, all the 11 ANNs that scored an average testing R^2^ > 0.80 have u_∗_in their models, indicating that information on atmospheric turbulence is important to incorporate in ET partitioning prediction if available. It may not be surprising that at these flooded sites E is mainly explained by atmospheric conditions such as VPD, T_air_, and turbulence (u_∗_) underlining their importance in the ANN partitioning routine. At sites with different surface and vegetation characteristics, such as dryland sites, it would be important to investigate the importance of other variables such as soil moisture, soil temperature, or leaf wetness. It would be expected that these, together with other energy balance components such as radiation, would play a larger role in explaining E at water limited sites.

### 4.2 Artificial Neural Network Validation Against Post-Flooding Periods and Leaf-Level Data

The validation of our models against data collected right after flooding (for East End, Mayberry Farms, and Sherman Island) and with leaf-level data (for East End only) indicated that models with less input variables (F11) performed better in comparison to the model that incorporated all 10 studied variables (F36). It might be that overfitting occurred when incorporating input variables that deal directly and/or indirectly with the same property/ factor (i.e., carbon assimilation). In this case, F36 includes er_Reichstein, wc_gf and GCC which are all related to carbon uptake by vegetation. Thus, even with a smaller average testing R^2^ value, models with fewer input variables (e.g., F11) still performed better than F36 during validation with ground-truth leaf-level and flooding data. Specifically, the ANN F11, which showed the best performance for all three of the sites with flooding data validation (East End, Mayberry Farms, and Sherman Island) included datetime + H_gf + VPD + T _air_ + u_∗_. The validation based on data collected right after flooding also emphasized the importance of validating the ANN partitioning routine against data collected during daytime periods. Some of the tested input variables showed strong differences in daytime and nighttime behavior (e.g., R_net_). Using these variables as inputs can lead to incorrect predictions for the nighttime-based ANN routine as seen in the poor performance of F15 for the flooding validation at East End and Mayberry Farms, despite a high testing R^2^ of 0.75 (Supplementary Table 1).

The flooding validation also highlights site-specific differences in the input variables that provided good predictions. While the best performance was achieved with the same model (F11) across all three validation sites, the behavior of the other tested models varied across sites. We recommend that the selection of input parameters for ANN partitioning of ET should be based on the unique site characteristics rather than a standardized set of variables since vegetation heterogeneity and other site level characteristics can influence ecosystem ET levels (Eichelmann et al., 2018).

This is also evident in the validation using data from the winter/senescent period, where F11 performed best at Mayberry Farms and Sherman Island, whereas F36 performed best at East End and West Pond. The overall performance of our ANNs in predicting E during the winter/senescent periods was also considerably lower in comparison to the flooding and leaf-level data validation. This is partially due to the smaller fluxes observed overall during this period, leading to larger relative errors. In addition, the assumption that all measured ET during the winter months represents solely E is likely incorrect. Especially at the sites with high vegetation cover (Mayberry and West Pond) it is likely that a small amount of T occurs during this time which would be included in the measured ET signal, leading to an apparent under-prediction of E for the ANN. For East End and Sherman Island, however, we can see that the ANNs are actually over-predicting E (Fig. 2), leading to consistent, albeit relatively small, negative T prediction in the winter months, specifically at East End (Fig. 4). It is unclear what is causing the discrepancy between measured and modeled E at East End and Sherman Island during the winter months. However, the fact that inclusion of variables linked to vegetation growth (GCC, wc_gf, er_Reichstein) reduced the over-prediction at both sites (e.g., F36 or F15) could indicate that E dynamics linked to phenology and vegetation cover are not adequately reproduced in models without these input variables at East End and Sherman Island.

Unfortunately, a limitation in our study is that we were not able to validate our results across all sites/sampling times due to a lack of leaf-level data collected from all sites, which is very time and labor intensive. In addition, no data were available from the initial flooding period at the West Pond wetland. Nonetheless, we are aware that validation of T/ET partitioning is quite scarce in the literature and that the data validated against our ANNs prove that good results can be achieved using the protocol tested here.

### 4.3 Artificial Neural Network Performance Across the Wetland Sites

Concerning the performance of all the 36 ANNs across the four wetlands analyzed in this study, West Pond showed smaller testing R^2^ values in comparison to the three other sites. Between-site differences reached up to 46% for the same model. The main reason for this divergence was likely the differing amounts of open water surfaces and density of the vegetation between these sites. West Pond, with little to no open water, is likely to see less E compared to the other wetlands (Eichelmann et al., 2018). In addition, West Pond also has the lowest water temperature and a very dense vegetation canopy decoupling the water surface from the atmosphere and leading to further reductions in E, especially at night (Drexler et al., 2004; Goulden et al., 2007; Eichelmann et al., 2018). Because our method predicts E based on nighttime data and calculates T based on the difference between total ET and E, if E values are small the relative accuracy of the prediction will decrease, which is reflected in the testing R^2^ values. However, because the E values are small, the absolute error of the predicted E and T would be proportionately small, hence the total T and E values can still be reliable. Unfortunately, we did not have a set of ground-truth validation data available for the West Pond site to investigate the true performance of the ANN ET partitioning. However, our comparison with the Scott and Biederman (2017) partitioned data and expected relationships based on the observed carbon fluxes and vegetation dynamics give us high confidence in the performance of the ANN partitioning routine at the West Pond wetland site. This shows that the ANN partitioning method can also be successfully applied in situations where nighttime E fluxes are small, indicating that it could be applicable to a large variety of ecosystems.

### 4.4 Comparisons with Other Partitioning Approaches

In comparison to other established methods in the literature our own approach using ANNs to determine the T/ET partitioning achieved very good results with fewer limitations, which makes it easier to apply in other contexts/ecosystems. For instance, Scott and Biederman’s (2017) method only works when there are enough years of data. The shortest dataset Scott and Biederman (2017) analyzed spanned eight years, which is a considerably long time period and reduces its applicability to shorter studies. Also, in the absence of climate consistency among sampling sites or if the research takes place in areas where fluxes are not limited by water availability (e.g., wetlands), their model fails to partition T/ET correctly, limiting it to relatively dry ecosystems. This was evident from direct comparisons with our own method, particularly for Sherman Island which has the shortest dataset (i.e., four years) and the highest area of open water, with the largest relative contribution of E (Fig. 4, 5).

Considering the partitioning methods proposed by Scanlon and Sahu (2008), Scanlon and Kustas (2010), and Skaggs et al. (2018), a priori knowledge on WUE and carbon uptake is required to apply their method. Consequently, the paucity of previous data/information or lack of equipment impede the application of this method to a broader audience. We tried to run the Scanlon and Kustas (2010) and Skaggs et al. (2018) partitioning methods for our wetland sites but were not able to retrieve reliable and meaningful partitioning results for any of the sites discussed in this study. We did not test the method proposed by Zhou et al. (2016) in this study, since we believe that some of the underlying assumptions are easily violated at the wetland sites investigated here. Most importantly, the Zhou et al. (2016) method is based on the assumption that some periods within the time series represent conditions without E and the water flux is entirely based on T (i.e., T = ET). This is most certainly not the case at flooded sites where we can reasonably expect that there will always be E, albeit in varying amounts. Additionally, the potential underlying WUE is assumed to be constant, which could be violated when multiple vegetation types or species are present, as is the case with our sites. Finally, virtually all the other methods discussed here lacked validation against ground-truth data in the original studies. We included several verification types for the ANN method in this paper, which gives us confidence that our approach using ANNs produces reliable and meaningful estimates for E and T in wetland ecosystems. The fact that our method does not rely on presumed relationships between water and carbon fluxes and was shown to work across a range of ecosystem properties from T to E dominated systems, provides an advantage against other methods that are limited to certain ecosystems or need specialized input data/equipment.

## 5 Summary

A novel T/ET partitioning method using Artificial Neural Networks (ANN) to predict daytime E from nighttime ET measurements in a combination with a range of environmental variables was presented and compared to previous methods from the literature. In comparison to other approaches, the ANN method achieved better results, particularly with shorter-term data (i.e., <5 years) and was successfully applied to flooded ecosystems. The order of variable inclusion (and importance) for the ANN construction was: vapor pressure deficit (VPD) > gap-filled sensible heat flux (H_gf) > air temperature (T_air_) > friction velocity (u_∗_) > other variables. The best performing ANN, model F11, used datetime, VPD, H_gf, T_air_, and u_∗_ inputs with an average testing R^2^ value across all sites of 0.85. This model also performed the best when validated against ground-truth leaf-level data and periods where sites were completely flooded with no T from vegetation. Our method sheds light on T/ET partitioning methods and applications. While here it has only been tested for flooded ecosystems, we present strong indicators that it could also perform well in other ecosystems, contributing to the understanding of the global water cycle.

## Acknowledgments

**Author contributions:** Conceptualization: E.E., D.D.B.; Data collection and processing: E.E., S.D.C., K.S.H., P.Y.O., D.S., A.V., J.V.; Formal analysis: E.E.; M.C.M; Writing—original draft: E.E., M.C.M; Writing—review and editing: D.D.B., A.V., K.S.H., P.Y.O.;

**Funding sources:** This work was supported by Enterprise Ireland (H2020-Proposal Preparation Support CS20202080); the California Department of Water Resources (DWR) through a contract from the California Department of Fish and Wildlife; and the United States Department of Agriculture (NIFA grant #2011-67003-30371). Funding for the AmeriFlux core sites was provided by the U.S. Department of Energy’s Office of Science (AmeriFlux contract #7079856). Dennis D. Baldocchi was supported by the McIntire-Stennis Capacity Grant funding to the California Agricultural Experiment Station. Kyle S. Hemes was supported by the California Sea Grant Delta Science Fellowship. This material is based upon work supported by the Delta Stewardship Council Delta Science Program under Grant No. 2271 and California Sea Grant College Program Project R/SF-70.

## Footnotes

1 Abbreviations: ANN = Artificial Neural Networks; EC = Eddy Covariance; E = evaporation; ET = evapotranspiration; GCC = vegetation greenness index; GEP = Gross Ecosystem Productivity; T = transpiration; WT = water table; WUE = Water Use Efficiency; VPD = vapor pressure deficit;

2 NB: There is some discussion in the community around the correct use of the terms evapotranspiration vs evaporation (Miralles et al, 2020). We have opted to follow the common use of the term evapotranspiration throughout this manuscript to describe the total biosphere-atmosphere water flux, including transpiration as well as direct evaporation from soil and surface waters.

