## Supplemental Table 2 for "A novel approach to partitioning evapotranspiration into evaporation and transpiration in flooded ecosystems"

**Supplementary Table 2** – Slope values of all 36 Artificial Neural Networks analyzed for evapotranspiration partitioning ordered by decreasing average slope across all four studied wetland sites (East End, Mayberry Farms, Sherman Islands, and West Pond) in the Sacramento-San Joaquin Delta, California, US.

| Model Name | Model Structure | East End | Mayberry Farms | Sherman Islands | West Pond | Average Slope |
|---|---|---|---|---|---|---|
| F36 | Mdate + H_gf + ustar + wc_gf + er_Reichstein + VPD + $T_{air}$ + GCC + Rnet + WT | 0.929 | 0.883 | 0.937 | 0.771 | 0.880 |
| F14 | Mdate + H_gf + ustar + VPD + $T_{air}$ + GCC + Rnet + WT | 0.913 | 0.863 | 0.938 | 0.758 | 0.868 |
| F20 | Mdate + H_gf + ustar + VPD + $T_{air}$ + GCC + WT | 0.921 | 0.858 | 0.933 | 0.701 | 0.853 |
| F35 | Mdate + H_gf + ustar + er_Reichstein + VPD + $T_{air}$ | 0.904 | 0.871 | 0.927 | 0.703 | 0.851 |
| F11 | Mdate + H_gf + ustar + VPD + $T_{air}$ | 0.877 | 0.852 | 0.927 | 0.669 | 0.831 |
| F1 | Mdate + ustar + VPD + $T_{air}$ + GCC + Rnet + WT | 0.883 | 0.833 | 0.893 | 0.704 | 0.828 |
| F19 | Mdate + ustar + VPD + $T_{air}$ + GCC + WT | 0.909 | 0.831 | 0.881 | 0.692 | 0.828 |
| F13 | Mdate + ustar + er_Reichstein + VPD + $T_{air}$ | 0.870 | 0.839 | 0.875 | 0.673 | 0.814 |
| F12 | Mdate + ustar + wc_gf + VPD + $T_{air}$ | 0.856 | 0.861 | 0.874 | 0.660 | 0.813 |
| F7 | Mdate + ustar + VPD + $T_{air}$ | 0.851 | 0.823 | 0.874 | 0.650 | 0.800 |
| F5 | Mdate + ustar + VPD | 0.857 | 0.812 | 0.855 | 0.621 | 0.786 |
| F16 | ustar + VPD + $T_{air}$ + GCC + Rnet + WT | 0.832 | 0.756 | 0.874 | 0.669 | 0.783 |
| F34 | Mdate + H_gf + VPD + $T_{air}$ | 0.805 | 0.745 | 0.818 | 0.567 | 0.734 |
| F33 | Mdate + H_gf + VPD | 0.808 | 0.736 | 0.813 | 0.545 | 0.726 |
| F2 | Mdate + ustar + $T_{air}$ | 0.773 | 0.735 | 0.821 | 0.456 | 0.696 |
| F32 | Mdate + VPD + $T_{air}$ + GCC + Rnet | 0.738 | 0.683 | 0.744 | 0.493 | 0.665 |
| F15 | Mdate + VPD + $T_{air}$ + GCC + Rnet + WT | 0.708 | 0.682 | 0.754 | 0.507 | 0.663 |
| F31 | Mdate + VPD + $T_{air}$ + GCC | 0.734 | 0.683 | 0.744 | 0.465 | 0.657 |
| F17 | Mdate + VPD + $T_{air}$ | 0.671 | 0.684 | 0.732 | 0.456 | 0.636 |
| F26 | Mdate + VPD | 0.703 | 0.660 | 0.710 | 0.432 | 0.626 |
| F8 | H_gf + VPD + $T_{air}$ | 0.554 | 0.620 | 0.756 | 0.485 | 0.604 |
| F28 | Mdate + H_gf | 0.743 | 0.613 | 0.619 | 0.385 | 0.590 |

| Model Name | Model Structure | East End | Mayberry Farms | Sherman Islands | West Pond | Average Slope |
|---|---|---|---|---|---|---|
| F6 | Mdate + ustar + GCC | 0.654 | 0.576 | 0.749 | 0.214 | 0.548 |
| F10 | er_Reichstein + VPD + $T_{air}$ | 0.471 | 0.561 | 0.681 | 0.413 | 0.532 |
| F4 | Mdate + ustar + Rnet | 0.635 | 0.587 | 0.707 | 0.193 | 0.531 |
| F22 | Mdate + $T_{air}$ | 0.621 | 0.562 | 0.637 | 0.299 | 0.530 |
| F3 | Mdate + ustar + WT | 0.597 | 0.565 | 0.694 | 0.199 | 0.514 |
| F18 | ustar + VPD + $T_{air}$ | 0.366 | 0.516 | 0.557 | 0.596 | 0.509 |
| F27 | Mdate + ustar | 0.625 | 0.555 | 0.679 | 0.175 | 0.509 |
| F9 | wc_gf + VPD + $T_{air}$ | 0.369 | 0.535 | 0.635 | 0.423 | 0.491 |
| F30 | Mdate + er_Reichstein | 0.582 | 0.449 | 0.592 | 0.142 | 0.441 |
| F25 | Mdate + Rnet | 0.565 | 0.501 | 0.548 | 0.125 | 0.435 |
| F23 | Mdate + GCC | 0.557 | 0.453 | 0.589 | 0.111 | 0.428 |
| F29 | Mdate + wc_gf | 0.570 | 0.451 | 0.548 | 0.141 | 0.428 |
| F24 | Mdate + WT | 0.513 | 0.446 | 0.510 | 0.108 | 0.394 |
| F21 | Mdate | 0.516 | 0.456 | 0.514 | 0.084 | 0.393 |
