## Supplemental Table 1 for "A novel approach to partitioning evapotranspiration into evaporation and transpiration in flooded ecosystems"

**Supplementary Table 1** – Testing $R^2$ values of all 36 Artificial Neural Networks analysed for evapotranspiration partitioning ordered by decreasing average $R^2$ across all four studied wetland sites (East End, Mayberry Farms, Sherman Islands, and West Pond) in the Sacramento-San Joaquin Delta, California, US.

| Model Name | Model Structure | East End | Mayberry Farms | Sherman Islands | West Pond | Average $R^2$ |
|---|---|---|---|---|---|---|
| F36 | Mdate + H_gf + ustar + wc_gf + er_Reichstein + VPD + $T_{air}$ + GCC + Rnet + WT | 0.935 | 0.900 | 0.925 | 0.804 | 0.891 |
| F14 | Mdate + H_gf + ustar + VPD + $T_{air}$ + GCC + Rnet + WT | 0.917 | 0.879 | 0.922 | 0.789 | 0.877 |
| F20 | Mdate + H_gf + ustar + VPD + $T_{air}$ + GCC + WT | 0.916 | 0.878 | 0.919 | 0.749 | 0.866 |
| F35 | Mdate + H_gf + ustar + er_Reichstein + VPD + $T_{air}$ | 0.908 | 0.885 | 0.913 | 0.747 | 0.863 |
| F11 | Mdate + H_gf + ustar + VPD + $T_{air}$ | 0.898 | 0.874 | 0.910 | 0.728 | 0.853 |
| F1 | Mdate + ustar + VPD + $T_{air}$ + GCC + Rnet + WT | 0.893 | 0.857 | 0.884 | 0.732 | 0.842 |
| F12 | Mdate + ustar + wc_gf + VPD + $T_{air}$ | 0.887 | 0.880 | 0.866 | 0.721 | 0.839 |
| F19 | Mdate + ustar + VPD + $T_{air}$ + GCC + WT | 0.896 | 0.853 | 0.873 | 0.729 | 0.838 |
| F13 | Mdate + ustar + er_Reichstein + VPD + $T_{air}$ | 0.882 | 0.854 | 0.870 | 0.723 | 0.832 |
| F7 | Mdate + ustar + VPD + $T_{air}$ | 0.872 | 0.843 | 0.864 | 0.709 | 0.822 |
| F5 | Mdate + ustar + VPD | 0.868 | 0.831 | 0.849 | 0.668 | 0.804 |
| F16 | ustar + VPD + $T_{air}$ + GCC + Rnet + WT | 0.861 | 0.724 | 0.865 | 0.709 | 0.790 |
| F34 | Mdate + H_gf + VPD + $T_{air}$ | 0.814 | 0.783 | 0.800 | 0.652 | 0.762 |
| F33 | Mdate + H_gf + VPD | 0.805 | 0.773 | 0.793 | 0.639 | 0.753 |
| F2 | Mdate + ustar + $T_{air}$ | 0.807 | 0.772 | 0.821 | 0.519 | 0.730 |
| F15 | Mdate + VPD + $T_{air}$ + GCC + Rnet + WT | 0.747 | 0.735 | 0.741 | 0.552 | 0.694 |
| F32 | Mdate + VPD + $T_{air}$ + GCC + Rnet | 0.740 | 0.725 | 0.740 | 0.552 | 0.689 |
| F31 | Mdate + VPD + $T_{air}$ + GCC | 0.742 | 0.724 | 0.735 | 0.542 | 0.686 |
| F17 | Mdate + VPD + $T_{air}$ | 0.713 | 0.724 | 0.717 | 0.533 | 0.672 |
| F26 | Mdate + VPD | 0.709 | 0.687 | 0.705 | 0.489 | 0.648 |
| F28 | Mdate + H_gf | 0.688 | 0.645 | 0.613 | 0.535 | 0.620 |
| F8 | H_gf + VPD + $T_{air}$ | 0.593 | 0.540 | 0.720 | 0.595 | 0.612 |

| Model Name | Model Structure | East End | Mayberry Farms | Sherman Islands | West Pond | Average $R^2$ |
|---|---|---|---|---|---|---|
| F6 | Mdate + ustar + GCC | 0.699 | 0.624 | 0.737 | 0.275 | 0.584 |
| F4 | Mdate + ustar + Rnet | 0.682 | 0.630 | 0.696 | 0.263 | 0.568 |
| F10 | er_Reichstein + VPD + $T_{air}$ | 0.507 | 0.595 | 0.681 | 0.477 | 0.565 |
| F22 | Mdate + $T_{air}$ | 0.643 | 0.613 | 0.639 | 0.364 | 0.565 |
| F3 | Mdate + ustar + WT | 0.651 | 0.598 | 0.680 | 0.266 | 0.549 |
| F27 | Mdate + ustar | 0.638 | 0.580 | 0.664 | 0.243 | 0.531 |
| F18 | ustar + VPD + $T_{air}$ | 0.412 | 0.460 | 0.566 | 0.647 | 0.521 |
| F9 | wc_gf + VPD + $T_{air}$ | 0.386 | 0.552 | 0.631 | 0.492 | 0.515 |
| F30 | Mdate + er_Reichstein | 0.598 | 0.549 | 0.598 | 0.196 | 0.485 |
| F29 | Mdate + wc_gf | 0.594 | 0.560 | 0.558 | 0.158 | 0.468 |
| F23 | Mdate + GCC | 0.582 | 0.514 | 0.594 | 0.161 | 0.463 |
| F25 | Mdate + Rnet | 0.554 | 0.540 | 0.546 | 0.185 | 0.456 |
| F24 | Mdate + WT | 0.528 | 0.487 | 0.519 | 0.151 | 0.421 |
| F21 | Mdate | 0.512 | 0.472 | 0.523 | 0.132 | 0.410 |
